## Supplementary Table for "Leveraging information between multiple population groups and traits improves fine-mapping resolution"

**Supplementary Tables**

| **Method** | **330-variant region** | **1610-variant region** |
| --- | --- | --- |
| **MGflashfm** | 106.2 (91.9, 120.8) | 684.3 (589.9, 717.3) |
| **MGfm** | 59.8 (59.1, 58.0) | 394.6 (380.6, 402.9) |
| **PAINTOR** | 1664.4 (757.4, 3824.9) | 1753.1 (1262.8, 2864.9) |
| **msCAVIAR** | 1274.7 (1253.5, 1294.1) | > 10 hours |

**Supplementary Table 1: Computational time of multi-group methods with varying region size.**
The median running times (with second and third quartiles) are given in seconds. Running time was measured over 100 replications in simulations of two traits in two groups. The traits had correlation 0.4 and sample sizes were 90,000 (EUR) and 10,000 (AFR). The *APOE* region chr19:45000000-45800000 (GRCh37/hg19), consisting of at most 1610 variants or a subset of at most 330 variants in a group, was used for all simulations.

| **Number of groups** | **2** | **3** | **4** | **5** |
| --- | --- | --- | --- | --- |
| **MGflashfm** | 684.3  (589.9, 717.3) | 761.7 (686.1, 842.7) | 913.3 (831.9, 1015.3) | 1089.5 (976.5, 1367.2) |
| **MGfm** | 394.6  (380.6, 402.9) | 452.9 (431.4, 467.5) | 551.3 (533.7, 572.9) | 622.7 (591.8, 890.0) |

**Supplementary Table 2: Varying number of groups, median MGflashfm and MGfm running times (with second and third quartiles).** Median time was measured over 100 replications in simulations of 2 traits in 2-5 groups. Traits had correlation 0.4 and sample sizes ranged from 10,000 to 90,000 among groups. The *APOE* region chr19:45000000-45800000 (GRCh37/hg19), consisting of at most 1610 variants in a group, was used for all simulations.

| **Number of Traits** | **2** | **3** | **4** |
| --- | --- | --- | --- |
| **Computational Time (min)** | 684.3  (589.9, 717.3) | 1024.9 (920.1, 1190.3) | 1214.2 (1111.6, 1337.9) |

**Supplementary Table 3: Computational time for MGflashfm with varying number of traits.** The median MGflashfm running time (with second and third quartiles) is given in seconds. Running time was measured over 300 replications in simulations of 2, 3, and 4 traits in two groups. The traits had correlation 0.4 and sample sizes were 90,000 (EUR) and 10,000 (AFR). The *APOE* region chr19:45000000-45800000 (GRCh37/hg19), consisting of at most 1610 variants in a group, was used for all simulations. As MGfm is for single traits, its speed is not considered here.

| **Method** | **T1: Pr(at least 1 cv in CS99)** | **T1: Pr(both cvs in CS99)** | **T2: Pr(at least 1 cv in CS99)** | **T2: Pr(both cvs in CS99)** |
| --- | --- | --- | --- | --- |
| **MGflashfm** | 1.0 | 0.960  (0.933, 0.987) | 1.0 | 0.970 (0.946, 0.994) |
| **MGfm** | 1.0 | 0.955 (0.926, 0.984) | 1.0 | 0.960 (0.933, 0.987) |
| **flashfm-AFR** | 1.0 | 0.965 (0.940, 0.990) | 1.0 | 0.970 (0.946, 0.994) |
| **flashfm-EUR** | 0.995 (0.985,1.0) | 0 | 1.0 | 0 |

**Supplementary Table 4: Calibration is retained by MGflashfm and MGfm upon exclusion of a causal variant in a group.** Two traits, each with two causal variants, were simulated in two groups. The causal variants for trait 1 are labeled as A, D and those for trait 2 are A, C, to indicate that one causal variant is shared between the traits. The A variant has MAF<0.01 in EUR and MAF>0.01 in AFR, and this is the variant that is removed from the EUR group. There are 200 replications.

**Supplementary Figures**

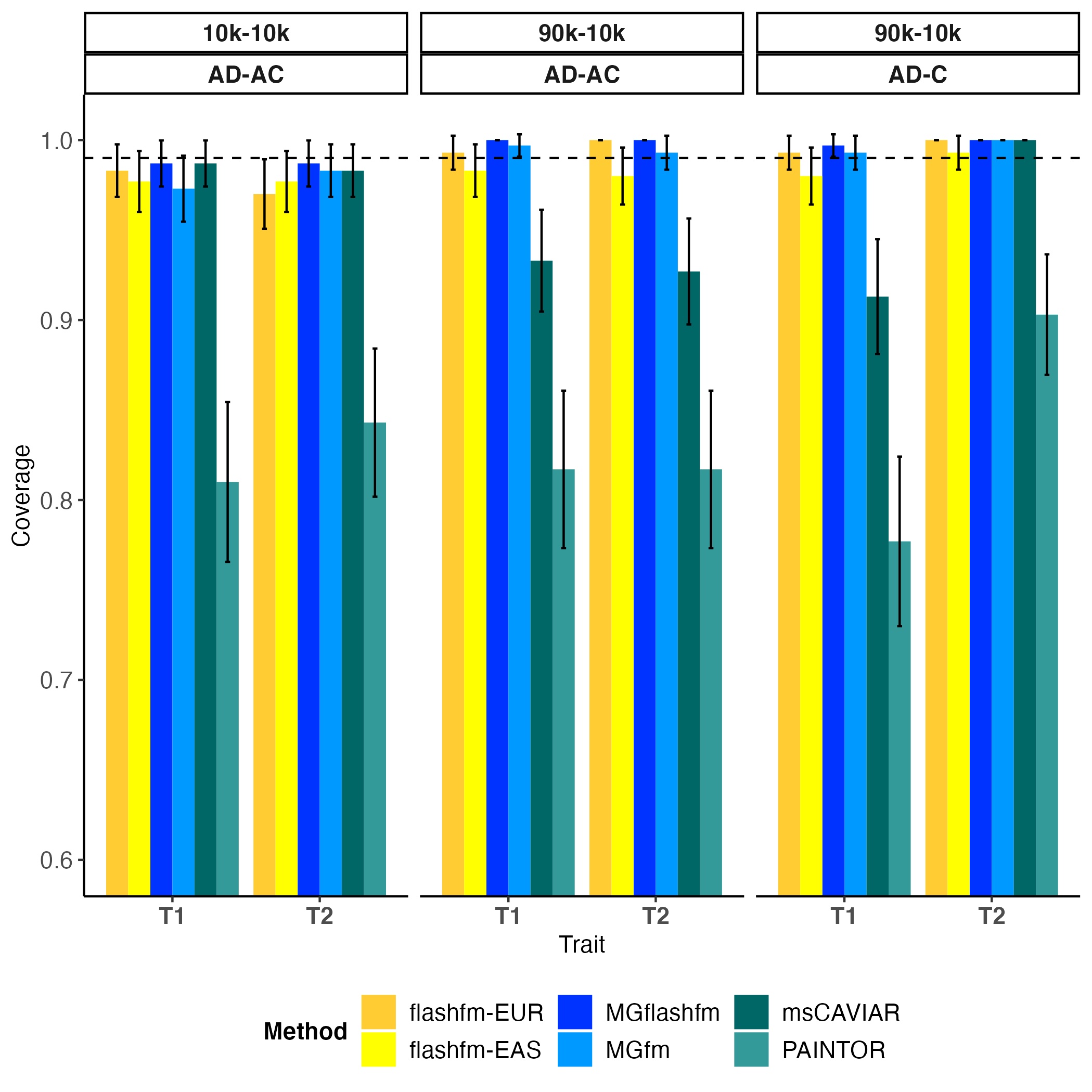

**Supplementary Figure 1. Flashfm, MGflashfm and MGfm, are well-calibrated in EUR-EAS fine-mapping.**
Coverage is measured as the probability that all causal variants are captured by the 99% credible set, estimated over 300 replications; 95% confidence intervals are shown around each coverage estimate. Flashfm-EUR and flashfm-EAS are multi-trait (single-group) fine-mapping for the indicated group and are well-calibrated in all settings, as are MGflashfm and MGfm. PAINTOR is not well-calibrated for all settings, while msCAVIAR is not well-calibrated for unequal sample sizes and for multiple causal variant settings. Within each panel the three simulation settings are shown as either having equal sample sizes of 10k each or sample sizes of 90k EUR and 10k EAS, and either two causal variants for each trait with one shared (trait 1: AD, trait 2: AC) or non-overlapping causal variants and one trait having a single causal variant (trait 1: AD, trait 2: C); any pair of causal variants have r^2^<0.5.

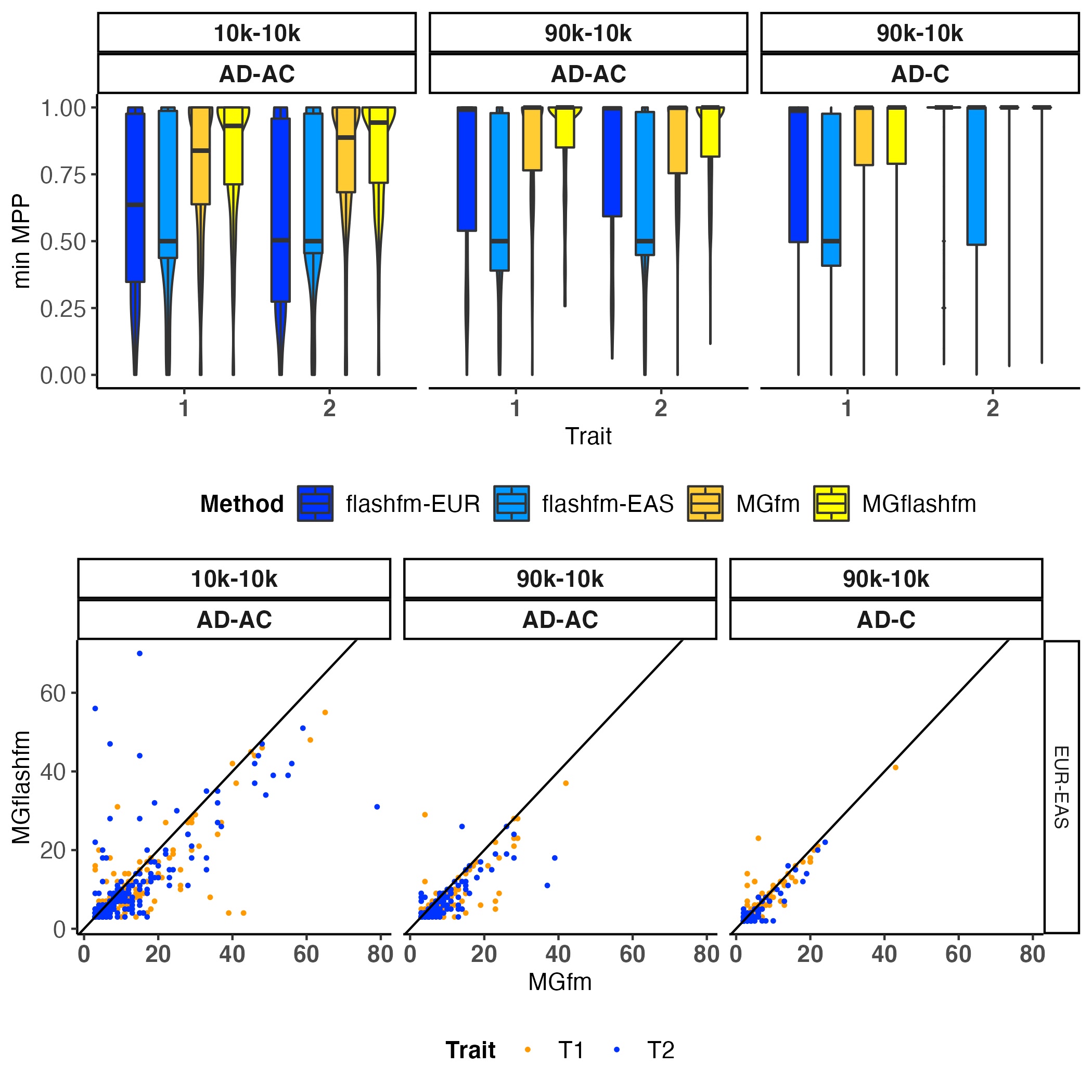

**Supplementary Figure 2. MGflashfm has the highest accuracy and resolution among calibrated methods for two traits in two groups.** For EUR-EAS simulations, three simulation settings are shown as either having equal sample sizes of 10k each or sample sizes of 90k EUR and 10k EAS, and either two causal variants for each trait with one shared (trait 1: AD, trait 2: AC) or non-overlapping causal variants and one trait having a single causal variant (trait 1: AD, trait 2: C); any pair of causal variants have r^2^<0.5 and there are 300 replications within each setting. The **upper panel** shows the distribution of the minimum MPP of causal variants for each trait; the median is given by the centre line, upper and lower quartiles are the box limits, whiskers are at most 1.5x interquartile range, and width indicates the frequency. This indicates that MGflashfm is best at prioritising causal variants when the traits share a causal variant or similar performance to MGfm when no sharing. The **lower panel** compares the sizes of 99% credible sets from MGflashfm and MGfm. This suggests that MGflashfm tends to have better resolution than MGfm.

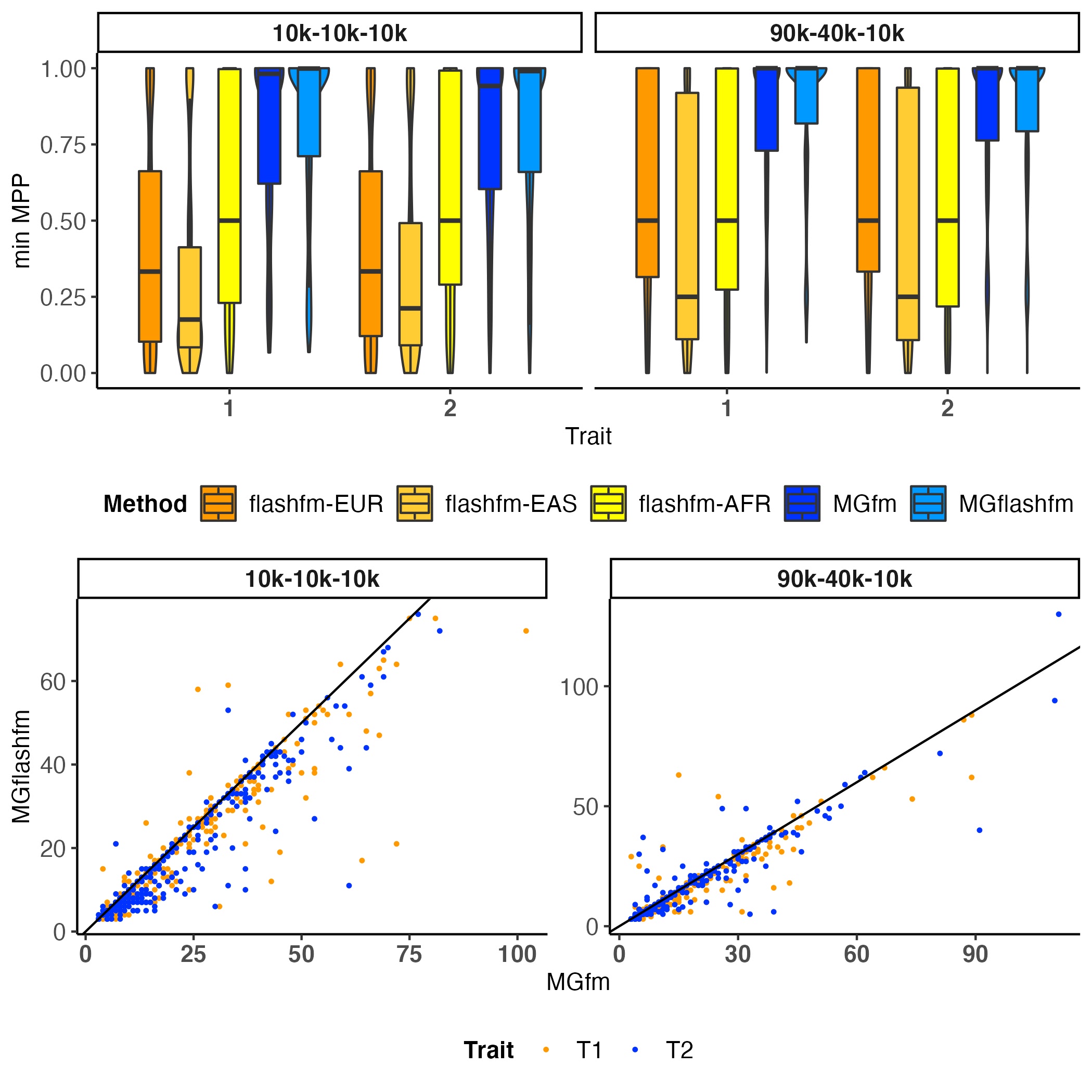

**Supplementary Figure 3. MGflashfm has the highest accuracy and resolution among calibrated methods for two traits in three groups.** For EUR-EAS-AFR simulations, two simulation settings are shown as either having equal sample sizes of 10k each or sample sizes of 90k EUR, 40k EAS, and 10k AFR. There are two causal variants for each trait with one shared (trait 1: AD, trait 2: AC) and any pair of causal variants have r^2^<0.5; there are 300 replications within each setting. The **upper panel** shows the distribution of the minimum MPP of causal variants for each trait; the median is given by the centre line, upper and lower quartiles are the box limits, whiskers are at most 1.5x interquartile range, and width indicates the frequency. This indicates that MGflashfm is best at prioritising causal variants. The **lower panel** compares the sizes of 99% credible sets from MGflashfm and MGfm. This suggests that MGflashfm tends to have better resolution than MGfm.
