## Supplementary Material for "Leveraging information between multiple population groups and traits improves fine-mapping resolution"

### Supplementary Section 1: Multi-group model description

The MGflashfm framework builds on that of flashfm [1] for multi-trait fine-mapping, which leverages information between traits to improve precision when there are shared causal variant(s) between traits. Multi-group fine-mapping has potential to further improve precision of fine-mapping due to differences in linkage disequilibrium (LD) between diverse population groups. For simplicity, we first consider the context of single-trait fine-mapping, then extend to multi-group multi-trait fine-mapping.

We first focus on derivations for the two group setting, which easily extends to more than two groups. Assume that for a single quantitative trait, we have $N_j,\ j=1,2$ measurements of the trait within group $j$. Also, within each group, the traits are transformed to meet conditional normality and homogeneity assumptions, conditional on covariates. Later, as in flashfm, we relax this so that a subset of individuals may have missing measurements for some of the traits. Here, we find an expression for the multi-group ABF of causal variant models and show that the information from single group analyses could be used to evaluate the multi-group ABF.

To find expressions of the log(ABF) for each of the joint and marginal models we use the approximation based on the Bayesian information criterion (BIC) from the null and causal models ($\mathrm{BIC}_0$ and $\mathrm{BIC}_1$, respectively)[2]. The log(ABF) approximation $(\mathrm{BIC}_0-\mathrm{BIC}_1)/2$, is expressed in terms of log likelihoods as

$$\log(\mathrm{ABF}) \doteq l_1 - l_0 - m\log(N)/2, \tag{1}$$

where $m$ is the number of causal variants in the model and $l_1$ and $l_0$ are the log likelihoods of the causal and null models, evaluated at the maximum likelihood estimates. Let $N = N_1 + N_2$, and denote a trait 1 model of $m_1$ variants by $M_1$ and, likewise, $M_2$ is a $m_2$-variant model for trait 2.

Using (1) and the fact that the groups are independent, we find that the the joint Bayes factor for models $M_1$ and $M_2$ for groups 1 and 2, having $m_1$ and $m_2$ variants, respectively, is given by

$$BF^{(1,2)}_{M_1,M_2} = BF^{(1)}_{M_1} \times \left(\frac{N_1}{N}\right)^{\frac{m_1}{2}} \times BF^{(2)}_{M_2} \times \left(\frac{N_2}{N}\right)^{\frac{m_2}{2}} \quad (2)$$

A natural joint prior probability for a $m_1$-variant trait 1 model with a $m_2$-variant trait 2 model is $p^{(1,2)}_{m_1,m_2} = p_{m_1} p_{m_2}$, where $p_{m_1}$ and $p_{m_2}$ are prior probabilities of a $m_1$-variant trait 1 model and $m_2$-variant trait 2 model, respectively; this considers the full joint model search space. Assuming that the groups share at least one causal variant, we add the restriction that the joint prior is only non-zero when the models overlap.

In particular, denoting the size (number of variants) in a model $M^{(k)}$ for trait $k$ by $|M^{(k)}|$, we have

$$\begin{aligned}&\Pr(|M^{(1)}| = m_1, |M^{(2)}| = m_2)\\ &= \Pr(|M^{(1)}| = m_1, |M^{(2)}| = m_2, M^{(1)} \cap M^{(2)} \neq \varnothing) + \Pr(|M^{(1)}| = m_1, |M^{(2)}| = m_2, M^{(1)} \cap M^{(2)} = \varnothing).\end{aligned}$$

However, we set the prior to 0 when $M^{(1)} \cap M^{(2)} = \varnothing$, so we introduce a correction factor $\tau_{m_1,m_2}$ such that

$$\Pr(|M^{(1)}| = m_1, |M^{(2)}| = m_2) = \Pr(|M^{(1)}| = m_1, |M^{(2)}| = m_2, M^{(1)} \cap M^{(2)} \neq \varnothing)\tau_{m_1,m_2},$$

so that the total joint prior probability of a $m_1$-variant group 1 model with a $m_2$-variant group 2 model in the reduced search space is anchored to remain the same as in the full model search space.

Let $S = \{(i,j) : |M^{(1)}_i| = m_1, |M^{(2)}_j| = m_2\}$ and $n$ be the number of SNPs in the region, then, we find $\tau_{m_1,m_2}$ as follows

$$\begin{aligned}\sum_{(i,j)\in S} p_{m_1} p_{m_2} &= \sum_{(i,j)\in S} p_{m_1} p_{m_2} 1\{M^{(1)}_i \cap M^{(2)}_j \neq \varnothing\}\tau_{m_1,m_2}\\ \binom{n}{m_1}\binom{n}{m_2} p_{m_1} p_{m_2} &= \left[\binom{n}{m_1}\binom{n}{m_2} - \binom{n}{m_1}\binom{n-m_1}{m_2}\right] p_{m_1} p_{m_2} \tau_{m_1,m_2},\end{aligned}$$

so that

$$\tau_{m_1,m_2} = \frac{\binom{n}{m_2}}{\binom{n}{m_2} - \binom{n-m_1}{m_2}} \tag{3}$$

So the joint prior probability is

$$p^{(1,2)}_{m_1,m_2} = p_{m_1} p_{m_2} 1\{M^{(1)} \cap M^{(2)} \neq \varnothing\} \tau_{m_1,m_2}, \tag{4}$$

where $\tau_{m_1,m_2}$ is the correction factor (3)

It follows from (2) and (4) that the joint posterior probability $PP^{(1,2)}_{M_1,M_2}$ for a particular model configuration $\{M_1,M_2\}$ may be found from only the $PP^{(1)}_{M_1}$, $PP^{(2)}_{M_2}$ of each model within their respective single-group fine-mapping model PPs, as follows:

$$\begin{aligned} PP^{(1,2)}_{M_1,M_2} &= p_{m_1} p_{m_2} BF^{(1)}_{M_1} \left(\frac{N_1}{N}\right)^{\frac{m_1}{2}} BF^{(2)}_{M_2} \left(\frac{N_2}{N}\right)^{\frac{m_2}{2}} 1\{M^{(1)} \cap M^{(2)} \neq \varnothing\} \tau_{m_1,m_2} \\ &= PP^{(1)}_{M_1} \left(\frac{N_1}{N}\right)^{\frac{m_1}{2}} PP^{(2)}_{M_2} \left(\frac{N_2}{N}\right)^{\frac{m_2}{2}} 1\{M^{(1)} \cap M^{(2)} \neq \varnothing\} \tau_{m_1,m_2} \end{aligned} \tag{5}$$

Let $C$ be a set of variants that compose a a multi-group model. This encompasses all group 1 - group 2 models that share at least one variant and $C$ is the collection of all variants in these models. So, the multi-group PP for set C is given by

$$PP_C = \sum_{\substack{ij: M^{(1)}_i \cup M^{(2)}_j = C, \\ M^{(1)}_i \cap M^{(2)}_j \neq \varnothing}} PP^{(1,2)}_{ij} \tag{6}$$

Finally, for variant $s$, multi-group MPPs (marginal posterior probability - probability that the variant appears in a model) are found from

$$MPP_s = \sum_{c: s \in c} PP_C.$$

This framework allows us to first evaluate evidence for multi-variant models within each group, accounting for group-specific LD. Then, evaluate joint evidence for a particular model configuration $\{M_i, M_j\}$, of model $M_i$ for group 1 with model $M_j$ for group 2, having already accounted for LD within each group.

Now, assume that we have measurements of $M$ quantitative traits within each group. We consider the same framework as above, but instead of making use of the single-trait fine-mapping model PPs from each group, we consider the multi-trait fine-mapping model PPs from flashfm applied to multiple traits within each group/study. This multi-group multi-trait fine-mapping approach is summarised by the following steps:

1. For each group, consider only variants that are present for all traits in that group, but do not intersect variants over groups;

2. Single-trait fine-mapping of each trait, within each group to obtain model PPs for each trait in each group;

3. Multi-trait fine-mapping (flashfm) within each group to leverage information between traits and obtain trait-adjusted model PPs for each trait in each group;

4. For each trait, use the above-described framework with $PP^{(1)}_{M_1}$ and $PP^{(2)}_{M_2}$ as found from multi-trait fine-mapping in groups 1 and 2, respectively.

This framework is extended to 3 groups, using similar arguments to the detailed 2-group setting previously described. Let $N = N_1 + N_2 + N_3$ and we have

$$BF^{(1,2,3)}_{M_1,M_2,M_3} = BF^{(1)}_{M_1} \times \left(\frac{N_1}{N}\right)^{\frac{m_1}{2}} \times BF^{(2)}_{M_2} \times \left(\frac{N_2}{N}\right)^{\frac{m_2}{2}} \times BF^{(3)}_{M_3} \times \left(\frac{N_3}{N}\right)^{\frac{m_3}{2}} \tag{7}$$

We assume that there is a shared causal variant between at least two of the three group models, which leads to a correction factor of

$$\tau_{m_1,m_2,m_3} = \frac{\binom{n}{m_2}\binom{n}{m_3}}{\binom{n}{m_2}\binom{n}{m_3} - \binom{n-m_1}{m_2}\binom{n-m_1-m_2}{m_3}};\ m_1 \geq m_2 \geq m_3 \geq 0.$$

So that the multi-group PP for a particular model configuration is

$$PP^{(1,2,3)}_{M_1,M_2,M_3} = PP^{(1)}_{M_1}\left(\frac{N_1}{N}\right)^{\frac{m_1}{2}} PP^{(2)}_{M_2}\left(\frac{N_2}{N}\right)^{\frac{m_2}{2}} PP^{(3)}_{M_3}\left(\frac{N_3}{N}\right)^{\frac{m_3}{2}} 1\{M_1 \cap M_2\ \neq \varnothing \text{ or } M_1 \cap M_3\ \neq \varnothing \text{ or } M_2 \cap M_3\ \neq \varnothing\}\tau_{m_1,m_2,m_3}$$

and the multi-group PP for a set C of variants is given by

$$PP_C = \sum_{\substack{h,i,j: M_h^{(1)} \cup M_i^{(2)} \cup M_j^{(3)} = C, \\ (h,i,j) \in S}} PP_{h,i,j}^{(1,2,3)},$$

where $S = \{(h,i,j) : M_h^{(1)} \cap M_i^{(2)} \neq \varnothing$ or $M_h^{(1)} \cap M_j^{(3)} \neq \varnothing$ or $M_i^{(2)} \cap M_j^{(3)} \neq \varnothing\}$
